# Identifying stationary phases in multivariate time-series for highlighting behavioural modes and home range settlements

**DOI:** 10.1101/444794

**Authors:** Rémi Patin, Marie-Pierre Étienne, Émilie Lebarbier, Simon Chamaillé-Jammes, Simon Benhamou

## Abstract

1. Recent advances in bio-logging open promising perspectives in the study animal movements at numerous scales. It is now possible to record time-series of animal locations and ancillary data (e.g. activity level derived from on-board accelerometers) over extended areas and long durations with a high spatial and temporal resolution. Such time-series are often piecewise stationary, as the animal may alternate between different stationary phases (i.e. characterised by a specific mean and variance of some key parameter for limited periods). Identifying when these phases start and end is a critical first step to understand the dynamics of the underlying movement processes.
2. We introduce a new segmentation-clustering method we called segclust2d. It can segment bi-(or more generally multi-) variate time-series and possibly cluster the various segments obtained, corresponding to phases assumed to be stationary. It is easy to use, as it only requires specifying the minimum length of a segment (to prevent over-segmentation) based on biological considerations.
3. Although this method can be applied to time-series of any nature, we focus here on two-dimensional piecewise time-series whose phases correspond at small scale to the expressions of different behavioural modes such as transit, feeding and resting, as characterised by two joint metrics such as speed and turning angles or, at larger scale, to temporary home ranges, characterised by stationary distributions of bivariate coordinates.
4. Using computer simulations, we show that segcust2d can rival and even outperform previous, more complex methods, which were specifically developed to highlight changes in movement modes or home range shifts (based on Hidden Markov or Ornstein-Uhlenbeck modelling, respectively), which, contrary to our method, require truly informative initial guesses to be efficient. Furthermore we demonstrate it on actual examples involving a zebra’s small scale movements and an elephant’s large scale movements, to illustrate the identification of various movement modes and of home range shifts, respectively.

## 1 Introduction

Landscapes are spatially and temporally variable at various scales (Levin, 1992), and animals are expected to adjust their movements to the characteristics of their local environment, so as to maximize the time spent in profitable (or safe) habitats and minimize time in adverse ones (Pyke, 1978). Recent advances in bio-logging have made it possible to acquire time-series of animal’s locations, and possibly ancillary data such as activity level derived from on-board accelerometers, over extended areas and long durations with high spatial and temporal resolutions. Such locational time-series, and the other ones that can be derived from them to describe the movement behaviour (e.g. turning angle, speed), are therefore expected to be piecewise stationary, i.e. to present a specific mean and variance for limited periods corresponding to stationary phases, alternating with rapid transition phases corresponding to changes of area or behaviour. Identifying these phases is a prerequisite to determine the biologically relevant scales of movement (Benhamou, 2014). It is therefore of paramount importance in two types of movement studies:

### Identifying behavioural modes

Foragers are generally expected to alternate intensive (area-concentrated) searching mode, characterized by high tortuosity and low speed, and extensive searching (transit) mode, characterized by low tortuosity and high speed (see Dias et al. 2009 for contrasting examples). This alternation of searching modes therefore results in piecewise “behavioural stationarity” when considering time-series of tortuosity and speed. Although different segmentation approaches have been developed to identify behavioural modes by looking at breakpoints (i.e. rapid transitions between stationary phases Barraquand and Benhamou, 2008; Gurarie et al., 2009; Nams, 2014), a more sophisticated approach based on Hidden Markov Models (HMM) has gained momentum in recent years. In this approach, the joint step lengths and turning angles calculated from successive relocations are categorized among a predefined number of different modes modelled as hidden states (Morales et al., 2004; Beyer et al., 2013; Langrock et al., 2012; McClintock et al., 2012; Michelot et al., 2016). However, the convergence of HMM may require specifying informative initial state-dependent probability distribution parameters (i.e. informative priors when HMM are designed in a Bayesian context), which can be difficult. Here, we aim at developing an alternative approach which does not require such a pre-specification.

### Identifying home range shifts

The recently emerging question of piecewise “locational stationarity” at the home range scale has been addressed in terms of movement scales (Benhamou, 2014), migration characteristics (Naidoo et al., 2012; Cagnacci et al., 2016) and of within-season shifts (Couriot et al., 2018). Indeed, for an animal that exploits various temporary home ranges, the time-series of relocations coordinates can be assumed to be stationary for a relatively long time (when the animal exploited the area where it established its temporary home range), then non-stationary for a relatively short time (when the animal left its home range until it established a new one), and so on. It is worth noting that a shift in home range does not necessarily involve a shift in mean location. It may also correspond to a change in variance if the animal enlarged or shrank its home range, e.g. due to a change in season (Naidoo et al., 2012; Monsarrat et al., 2013) or in reproductive status. Various methods have been proposed to detect home range shifts. The simple univariate approach based on the change of the beeline distance from a starting point (Bunnefeld et al., 2011) appears to be convenient in some cases but fully ignores movements leading the animal at a similar distance from the starting point but in another direction. A more complex approach rests on multi-state Ornstein-Uhlenbeck modelling (OUM Breed et al., 2017; Gurarie et al., 2017). However, as it requires that all home range phases and shifts are explicitly modelled, this approach tends to become cumbersome when there are several shifts to consider. Furthermore, it may require truly informative initial guesses to correctly detect small shifts. We therefore aim at developing an alternative approach that could be more efficient than an OUM-based approach to detect home range shifts and simpler to use. Additionally, as detecting shifts in behavioural modes and in home ranges settlements are conceptually similar, we focused on a generic approach that can be applied to both types of studies.

Here we introduce a new method, called segclust2d, able to segment a bi-(and more generally multi-) variate time-series, and to cluster similar segments (corresponding to stationary phases) in a common class (corresponding to a given state) if desired. We demonstrate that this method, which is easy to use, can successfully identify stationary phases corresponding to temporary home ranges when based on bivariate locational time-series, as well as movement modes when based on bivariate time-series of metrics such as speed and tortuosity. It thus offers an efficient and user-friendly alternative to previous, more complex, approaches. Furthermore, as this model applies to multivariate piecewise stationary time-series based on any kind of metrics, it can integrate additional time-series of ancillary data (e.g. activity level derived from on-board accelerometers) for a better segmentation-clustering of movement data.

## 2 Methods

### 2.1 Statistical model and parameter estimation

#### General principle

Consider a multivariate piecewise stationary time-series assumed to be regular (no gaps), composed of an unknown number of stationary phases. The *C* components of this signal (each corresponding to a univariate time-series) are assumed to be statistically independent (conditionally to the stationary phases) and should be normalized if they are of different nature, so as to have the same weight in subsequent procedures. One needs a reliable statistical model to find and characterize these phases and possibly to cluster them when they are assumed to be the expressions of a limited number of unobserved states of the underlying process (e.g. behavioural modes). Likelihood-based segmentation methods provide a suitable statistical framework to detect changes of phases but raise two main issues from a statistical and algorithmical point of view: (i) determining the optimal number of segments and (ii) for a given number of segments, finding the optimal segmentation, i.e. determining the locations of the starting/ending points of the segments (called breakpoints). The latter reduces to a well-known discrete optimization problem solved using a dynamic programming algorithm introduced by Bellman (1954; for a recent example, see Rigaill 2015). With *n* sites where the signal can be cut and *K* segments, the dynamic programming algorithm reaches the exact maximum likelihood solution with a complexity in *O*(*n*^2^*K*), drastically smaller than the complexity in *O*(*n^K^*) involved by a force brute algorithm when exploring the whole segmentation space. We will first introduce the models and the estimation procedure to optimally segment a multivariate signal for a predefined number of segments *K* and possibly (if clustering is required) a predefined number of states *M*. Afterwards, we will show how the optimal number of segments and possibly of clusters can be found based on a penalized likelihood criterion. Our approach is based on Lavielle (2005)’s segmentation method of univariate signals and its extension by Picard et al. (2007) to segment and cluster DNA sequences without assuming any kind of distribution for the segment lengths, such as a geometric distribution as HMM implicitly do (Karlin and Taylor, 1975).

#### Optimal segmentation in *K* segments, with optional clustering in *M* states

Assume that there are *K* stationary phases in a bivariate time-series with total length *n*. A stationary phase corresponds to a segment. It is defined by a sequence of consecutive random variables sharing the exact same distribution, in particular the same mean ***μ*** and variance matrix ***σ***^2^. As soon as one of these parameters changes, a new segment starts. The *C* components within a given segment *k* ∈ [1*, …, K*] starting at time *t* = *t_k−_*_1_ + 1 and ending at time *t* = *t_k_* (with *t*_0_ = 0 and *t_K_* = *n*) are modelled as sequences of Gaussian independent random variables *bY_t_*, for *t* = 1, …, n. In the absence of clustering (segmentation-only model), *bY_t_* (with *C* components labelled 1 to *C*) at time *t* is modelled simply as follows:

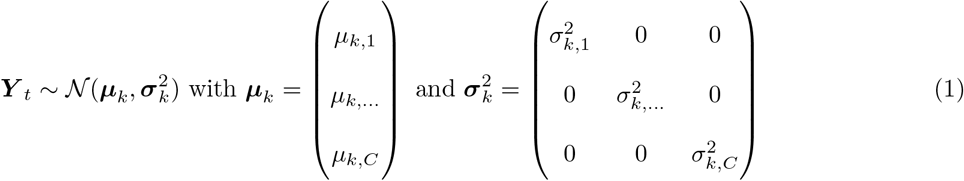

where ***μ****k* and 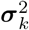 are the mean and variance matrix for segment *k*. As the model parameters to be estimated vary independently between segments, dynamic programming can be used to segment the multivariate signal at best in *K* segments. Its application is straightforward in this case, as it relies on the log-likelihood of each segment, which is simply equal to the sum of the log-likelihoods of the *C* components.

In the segmentation-clustering model, a state *m*, among *M* possible states, is assigned to every segment, and random variables within a segment classified in state *m* are all assumed to share the same mean ***μ***_*m*_ and the same variance 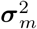, different from those involved in other states. More formally, let (*S_k_*), with *k* = 1, …, *K*, denote the state of the segment *k*: *S_k_* is a latent random variable taking values in [1, …, *M*]. It is modelled through a multinomial distribution of parameters *π* = (*π_m_*) with *m* = 1, …, *M*, where*π_m_* corresponds to the probability for a segment to belong to state *m*. Given *S_k_* = *m*, the value of the bivariate signal at time *t*, ***Y*** _*t*_, is therefore modelled as:

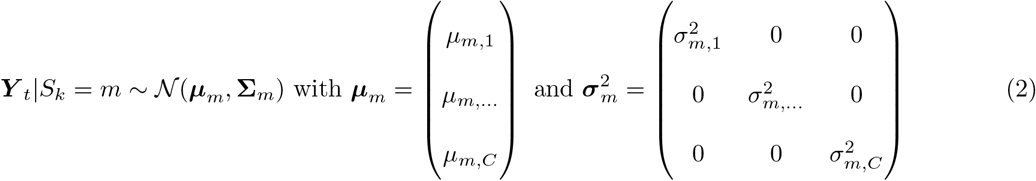

As parameters (*π_m_*, *μ_m_*, 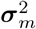) that characterise any state *m* are unknown and are to be estimated, resulting in a mixture distribution where segments are linked in terms of parameters, the optimal segmentation cannot anymore be obtained using dynamic programming alone. Following Picard et al. (2007), we designed the following two-step procedure, which is iterated up to convergence.

1. Given a set of parameters (*π_m_*, *μ_m_*, 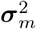) with *m* = 1, …, *M*, the best segmentation in *K* segments is obtained using dynamic programming.
2. Given this segmentation, the values of parameters are estimated using an expectation-maximization algorithm which is commonly used in latent variable modelling (Dempster et al., 1977).

By mixing dynamic programming and expectation-maximization through this iterative procedure, segmentation and clustering processes work jointly (rather than the latter after the former) leading to the optimal segmentation given *K* and *M*. However, an additional smoothing procedure is required to solve possible convergence issues (see details in Supporting information 1).

#### Finding the optimal numbers of segments and states

For both methods (segmentation-only and segmentation-clustering), a minimum segment length *L_min_* has to be set not only to speed up the algorithm, but also, more fundamentally, to prevent over-segmenting, based on biological considerations. For example, setting *L_min_* to a value of a few weeks when analysing locational time-series will prevent the algorithm from considering an area exploited only for a few days, corresponding to foray outside the usual home range or to stop-over during migration, as a distinct home range. Similarly, setting *L_min_* to a value of several minutes when looking for changes in behavioural modes will prevent force the algorithm to assign a given movement bout to a given behavioural mode only if this bout is long enough, even though it may be punctuated by ephemeral events related to another behaviour. To obtain the optimal solution, we calculated the likelihood of all number of segments 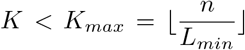, and for any number of states *M* (< *K*) one wishes to consider if clustering has to be involved. In this case, the optimal values of *K* and *M* are determined as those that maximize a Bayesian Information Criterion (BIC; Schwarz et al. 1978)-based penalised likelihood (Supporting information 1). However, as it will be shown in the Results section, it is usually preferable to consider a single value of *M* based on biologically relevant grounds than to let the model determine an optimal number of states based on a statistical basis (see also Pohle et al. 2017). When only segmentation is involved (no clustering), the optimal number of segments K is based in agreement with Lavielle (2005) on maximizing a K-penalized likelihood curve (Supporting Information 1).

### 2.2 Computer simulations

We run simulations to assess the ability of our approach to detect home range shifts and changes in behavioural modes from bivariate time-series, and to compare it with that of other (OUM-based and HMM-based) methods. For each set of parameters of each type of simulation, we simulated 100 replicates. Distances are expressed in arbitrary unit length *u*.

#### Home range shifts

For simplicity, the animal was assumed to behave as a central place forager. We simulated its fine-scale movement as a central-place biased correlated random walk, which results in a probability of presence decreasing exponentially with the distance D to the central place (Benhamou, 1989): at each time step, the animal turns by an angle *α_i_* drawn from a wrapped Gaussian distribution with a null mean and standard deviation 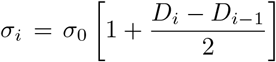, and progresses by 1 unit length (1*u*) in the new direction. In a batch of simulations, *σ*_0_ was set to 0.5 radians, and the central place was first set at a given location for the first 10,000 time steps (phase 1), then shifted to another location by 60*u* in both X and Y for 10,000 additional time steps (phase 2), resulting in disjoint home ranges, and then shifted to a third location by 20*u* in both X and Y for 10,000 additional time steps (phase 3), resulting in overlapping home ranges. In another batch of simulations, the central place remained at the same location for the 30,000 time steps, but *σ*_0_ was set to 0.7 radians for time steps 10,001 to 20,000 (phase 2) and to 0.5 radians otherwise (phases 1 and 3), involving a transitory enlargement of the home range. We finally sub-sampled the data sets by keeping one location every 60. The home range phases were then defined by 166 locations each, with low serial correlation, and were thus similar to actual datasets that are commonly used in home range studies. Note that in our approach that focuses on the contrast between the stationary phases, the actual lengths of these phases do not matter, provided they are longer than *L_min_*.

#### Changes in behavioural modes

We simulated a random search movement as a correlated random walk where three types of activity – immobility (resting or standing), intensive (area-concentrated) searching and extensive searching (transit) – alternate, each one lasting 20 time steps, this 60-step sub-series being repeated 5 times. The step lengths *L_i_* were drawn from a log-normal distribution with a mean equal to 0.5*u* in the intensive mode or 1.0*u* in the extensive mode, and with a standard deviation equal to 1/10th of the mean in both modes. Turning angles *α_i_* were drawn from a wrapped Gaussian distribution with a null mean and a standard deviation equal to 0.4 radians in the intensive mode or 0.3 radians in the extensive mode. To mimic possible factors (e.g. GPS recording noise) that can blur the contrast between the modes, the locations obtained in this way, as well those obtained for immobility phases, were submitted to bivariate Gaussian random noise with a null mean and various standard deviations *ζ*. Note that in order to assess the ability of a method to segment a behavioural time-series, the precise movement rules used in simulations are not important. What really matters is the contrast between the different phases: with a high contrast, all methods should work well, whereas with a low contrast, all methods should fail, whatever the movement rules considered. In the results section, we will present the results obtained with a standard deviation of the noise *ζ* set to 0.3, involving a moderate contrast between the three modes. The results obtained with a lower noise (*ζ* = 0.2; high contrast) or higher noise (*ζ* = 0.4; low contrast) are provided in Supporting Information.

### 2.3 Metrics

For identifying home range shifts, the two signal components considered are orthonormal Cartesian coordinates (*x_i_*, *y_i_*)) of locations (GPS locations expressed in decimal degrees as longitude and latitude therefore require to be transformed in terms of easting and northing through a classical projection such as UTM). For identifying behavioural modes, the two components usually considered in HMM-based approaches are the classical metrics corresponding to the step lengths *L_i_* and the turning angles *α_i_*, computed from locations recorded at constant time intervals ∆*t*, and therefore acting as proxies for linear (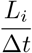) and angular (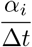) speeds, respectively. We used such metrics for comparative purpose, but we also tested some variants, assumed to improve the contrast between the different modes. We computed the linear speed as 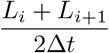. Although this basic smoothing introduces some serial correlation which is not taken into account in our model, it should result in a less noisy signal. Furthermore, angular speed may show faded changes between searching modes because the intensive mode usually involves both a decrease in linear speed and an increase in path tortuosity but angular speed mechanically increases with both of them (Benhamou and Bovet, 1989; Barraquand and Benhamou, 2008).

We therefore computed turning angles 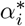 based on a constant step length *r* rather than at constant time interval. For this purpose, each location ***X***_*i*_ = (*X_i_, Y_i_*) is considered the centre of a virtual circle with radius *r*, and the entrance and exit locations *P_en_* and *P_ex_* are determined through linear interpolation (Appendix 2). The turning angle 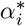 is then computed in [−*π*, *π*] as the angular deviation between vectors *P_en_* → *X_i_* and *X_i_* → *P_ex_* (both with length *r*) rather than vectors *X*_*i*−1_ → *X_i_* (with length *L_i_*) and *X_i_* → *X_i_*_+1_ (with length *L_i_*_+1_) as done to compute *α_i_*. When both *L_i_* and *L_i_*_+1_ are larger than *r*, one gets 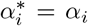, whereas 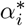 tends, on average, to be larger (random search paths) or smaller (directed paths) than *α_i_* when *r* is larger. We set *r* to the median of the step length distribution. In our simulations, we noticed that using a larger radius tends to improve the discrimination between the fast and slow movement modes but to worsen the discrimination between the slow movement mode and the immobility mode. We also tested the two orthogonal signals provided by the ‘persistence speed’ 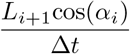 and ‘turning speed’ 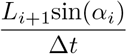 (Gurarie et al., 2009; Gloaguen et al., 2015).

### 2.4 Practical implementation of the method

Both segclust2d procedures (segmentation-only and segmentation-clustering) have been currently implemented to work on bivariate times-series (the only ones we considered in this paper) in a R package (https://cran.r-project.org/package=segclust2d), which optionally allows *K_max_* to be set to a value lower than 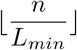 for preventing the algorithm from wasting time in looking for unlikely solutions. An integrated module makes it possible to derive the various movement variables mentioned in this paper from locations data. Because our approach requires large amounts of computer memory, it cannot deal with too long time-series (¿ 10000 values) on small desktop computers. To circumvent this limit, some sub-sampling is automatically performed when necessary. It is worth noting however that, even in absence of any memory constraints, it is usually not a good practice to attempt to directly segment very long series, which encompass both very large scale phenomena thanks to their large extent and very small scales phenomena thanks to their high resolution. Indeed, small-scale data are usually not relevant for analysing large-scale patterns and therefore act more as noise than as information in this context. Thus, in home range simulations, subsampling by a factor 60 makes it possible to dramatically shorten the time-series by eliminating fine-scale movements (klinokinetic process), which are characterised by a high level of serial correlation in location and in direction. Such details are clearly not relevant for the question of home range shifts, where only the overall phase-dependent mean and/or the variance of locations matter (accordingly, our approach ignores serial correlations occurring in any stationary phase). Conversely, for fine-scale movement studies, the characteristics of the environment are liable to change (e.g. due to seasonal variations) when considering a time-series running over an extended duration, possibly leading to change in the characteristics of the behavioural classes expected. It appears therefore preferable to consider the various phases (e.g. seasons) separately rather than to attempt to deal with the long time-series as a whole.

## 3 Results

### 3.1 Identifying home range shifts

#### Simulated movements

Figure 1 shows an example where the central place of the home range was shifted by 60 u between phases 1 and 2, and by 20*u* between phases 2 and 3, in both easting (X) and northing (Y), and an another example where the home range was enlarged during the phase 2 with respect to phases 1 and 3. The segclust2d/segmentation-only procedure (with Lmin set to 45 locations, corresponding to 2700 time steps because of the 1/60 subsampling) was able to correctly determine the true number of phases (3) in 98 out of the 100 replicates involving shifts in mean location (i.e. central place), and 88 out of the 100 replicates involving shifts in variance (i.e. change in home range size). In contrast, the OUM-based algorithm “marcher”, which was specifically developed by Gurarie et al. (2017) to identify home range shifts in mean location, requires that the number of shifts has been specified, and is unable to detect shifts in variance. When the true number of phases (3) has been specified, our approach was able to correctly estimate the occurrences of the various shifts (mean±SD = 10152±79 and 20092±188 time units for the 60*u* and 20*u* shifts in mean location, respectively; mean±SD = 10035±1184 and 19942±1314 time units for the first and second shifts in variance of the same amplitude). When the actual home range centres and shift dates were provided as truly informative initial guesses to Gurarie et al. (2017)’s method, it was also able to correctly detect the two shifts in mean location (mean±SD = 10152±82 and 19967±239 time units for the 60*u* and 20*u* shifts in mean location, respectively). In contrast, when no information was provided, Gurarie et al. (2017)’s method, which then relies on a k-means procedure to get initial guesses, was still able to detect the large shift in location (mean±SD = 10152±937 time units) but was unable to detect the small one (the shift occurred at random between 15000 and 30000 time units, based only on 75 replicates, as the algorithm failed to provide any result for 25 replicates).

**Figure 1:**
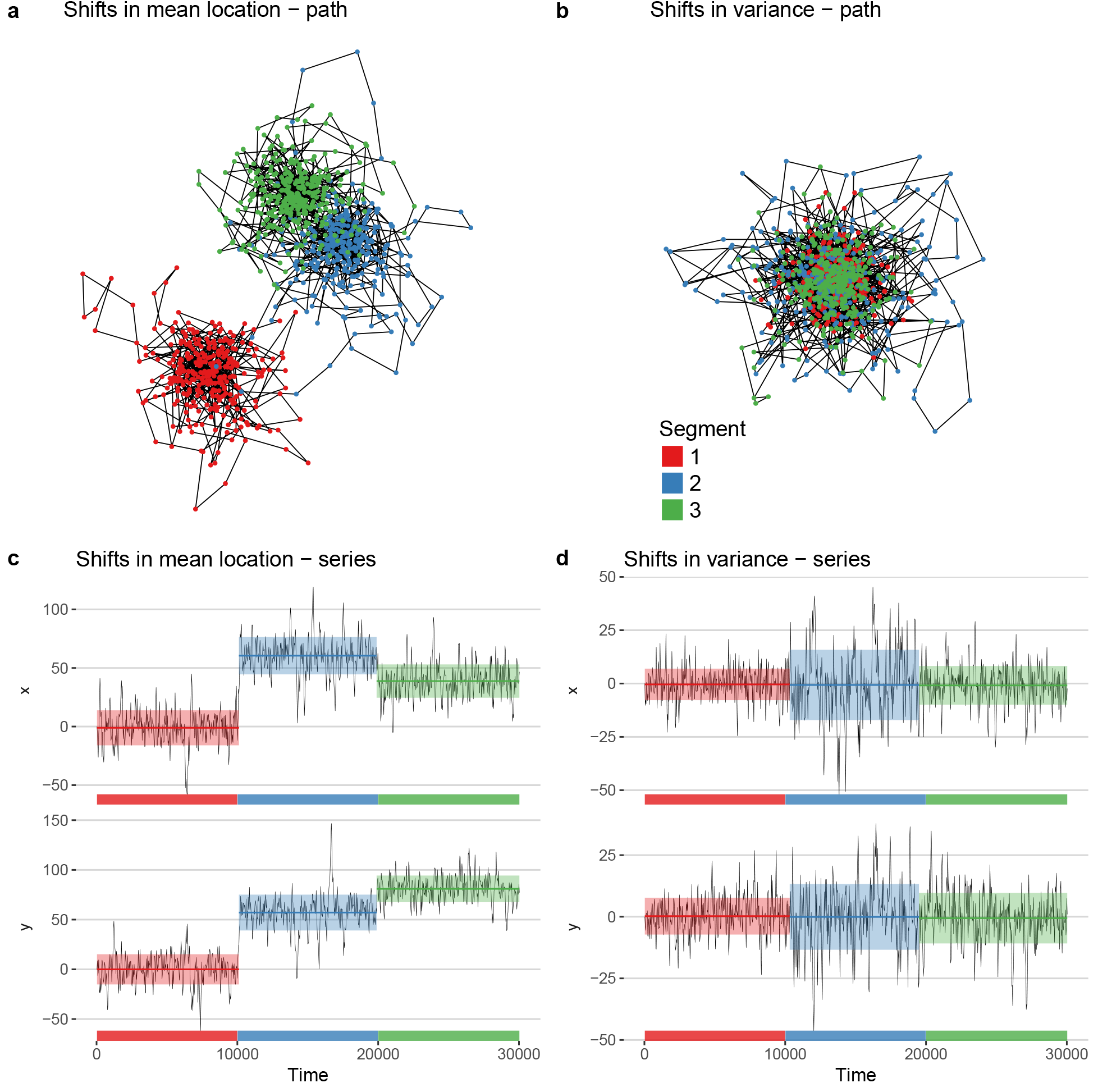
Examples of segmentation-only of simulated movements using segclust2d to high-light home range phases and shifts. Top panels show the simulated paths (after 1/60 subsampling) corresponding to three home range phases (two shifts), either in mean location (a) or in variance (b). The corresponding time series for both location coordinates (x, y) are presented in panel (c) and (d), respectively. The horizontal colour bars running along the time axis show the true occurrences of the three phases, whereas the coloured bands appearing over the x and y signals show their occurrences as estimated using the segclust2d/segmentation-only method (with Lmin = 45 locations, i.e. 2700 unit times) and provide the estimated mean (plain horizontal line running in the middle of the band) ± standard deviation (band width) for each segment separately.

#### Illustrative example

We used the GPS track of an African Elephant (*Loxodonta africana*), recorded for > 2.5 years to illustrate the way the segclust2d/segmentation-only procedure can identify home ranging phases and shifts (Fig. 2). Considering that a time-series is (roughly) stationary when the partial means and variances obtained for its first and second halves or for its three thirds are not markedly different, the whole time-series of easting and northing coordinates can be said to be both stationary, corresponding to a large multiannual (possibly lifetime) home range, and piecewise stationary, corresponding to temporary (possibly seasonal) smaller home ranges. It can therefore be segmented to highlight these temporary home ranges and shifts between them. However, some of the phases so highlighted are clearly nonstationary. In particular, segments 1 and 5 correspond mainly to a slow south-westwards migration (rather than a temporary home range) between the two core areas of the multiannual home range. Segment 2 also corresponds to a nonstationary, migratory (southwards moving) phase, which went through an area used as a temporary home range during segments 4 and 6. This indicates that a same area can be used in different ways at different periods.

**Figure 2:**
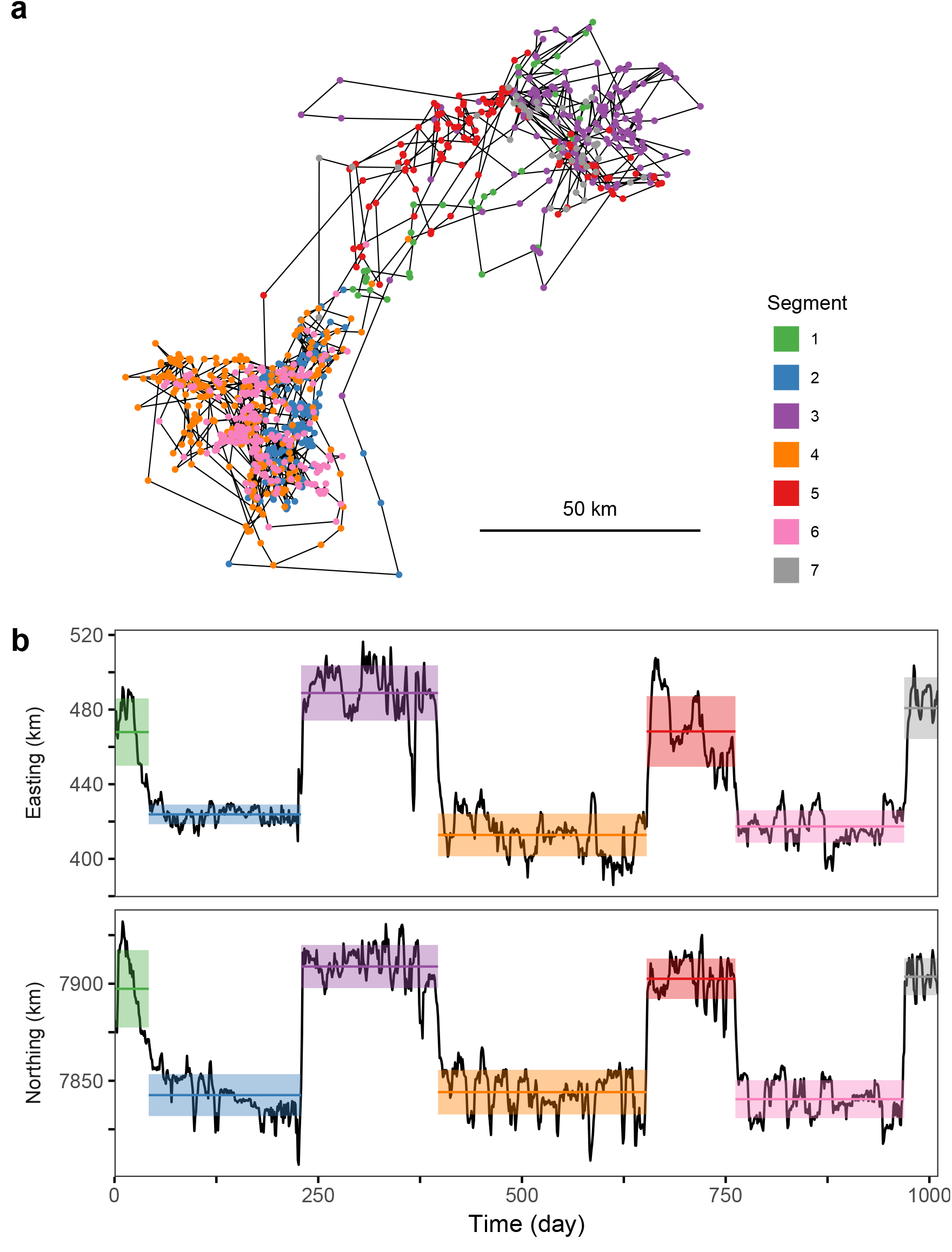
Example of segmentation-only of an African elephant’s movement recorded over 1000 days using segclust2d to highlight home range phases and shifts. (a) Rough path representation obtained by linking the locations subsampled so as to keep a single GPS location per day; (b) Corresponding time series of locations coordinates (easting and northing). The coloured bands appearing over the time series show the estimated mean (plain horizontal line running in the middle of the band) ± standard deviation (band width) of each of the seven segments obtained using the segmentation-only method with Lmin = 30 days.

### 3.2 Identifying behavioural modes

#### Simulated movements

An example of path with three behavioural modes (extensive searching, intensive searching and resting) is shown in Fig. 3 with the corresponding time-series in terms of turning angle 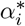 and smoothed speed 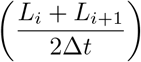. In this example, the segclust2d/segmentation-clustering procedure appears able to detect the true number of modes (*M* = 3) and to attribute almost all locations to the right mode. We compared our method with a HMM-based method specifically designed to deal with movement data (Michelot et al., 2016; McClintock and Michelot, 2018) when the true number of modes has been specified. The results obtained from 100 replicates showed that our procedure rivals with the HMM-based method although the latter was helped by initial state-dependent probability distribution parameters which were tuned to their true values for each behavioural state (Fig. 4 with medium noise level *ζ* = 0.3). With very low noise level (*ζ* =*<* 0.2), an excellent fit was obtained with all methods and metrics considered, whereas with a very high nose level (*ζ* => 0.4) the percentage of correct state assignment became closer to the value expected for a random assignment (33%; see Supporting information 3.1 results with *ζ* = 0.2 and *ζ* = 0.4). It also appeared that, as expected, the angular (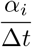) and linear (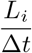) speeds are not the most suitable the metrics for detecting behavioural changes. Thus, better results were obtained with both methods when using any other of the couples of metrics considered. The best fits were obtained with turning angle 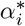 or it absolute value |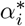| and smoothed speed. When the true number of modes is unknown, our method can also estimate this number as the most likely number of clusters, but the fraction of correct estimate is too low to consider the result as reliable (Supporting information 3.2).

**Figure 3:**
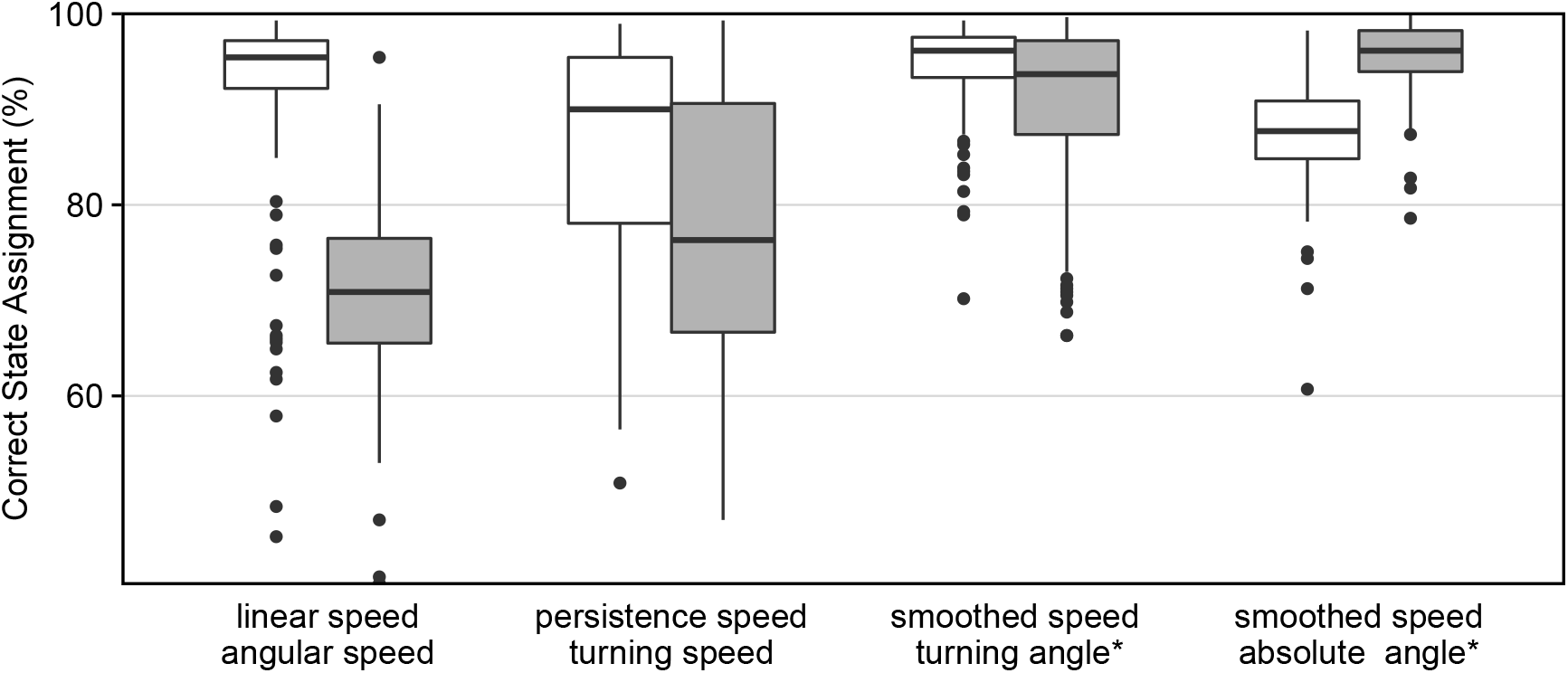
Comparative performances of segclust2d/segmentation-clustering vs. a HMM-based method for highlighting behavioural changes. The boxplots show the proportion of correct state assignments, obtained for various bivariate signals when the true number of states is known (M = 3), as estimated from 100 replicates simulated with the same parameters as to the one illustrated in fig. 4 (noise *ζ* = 0.3). The star (*) indicates turning angles computed with a constant step length, in terms of arithmetic (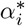) or absolute (—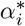—) values. The white boxplots show the results obtained with HMM-based R package momentuHMM (McClintock and Michelot, 2018), with Gaussian priors set to the true means and variances of the various metrics in the different states. The grey boxplots shows the results obtained using segclust2d/segmentation-clustering with *L_m_in* = 10.

**Figure 4:**
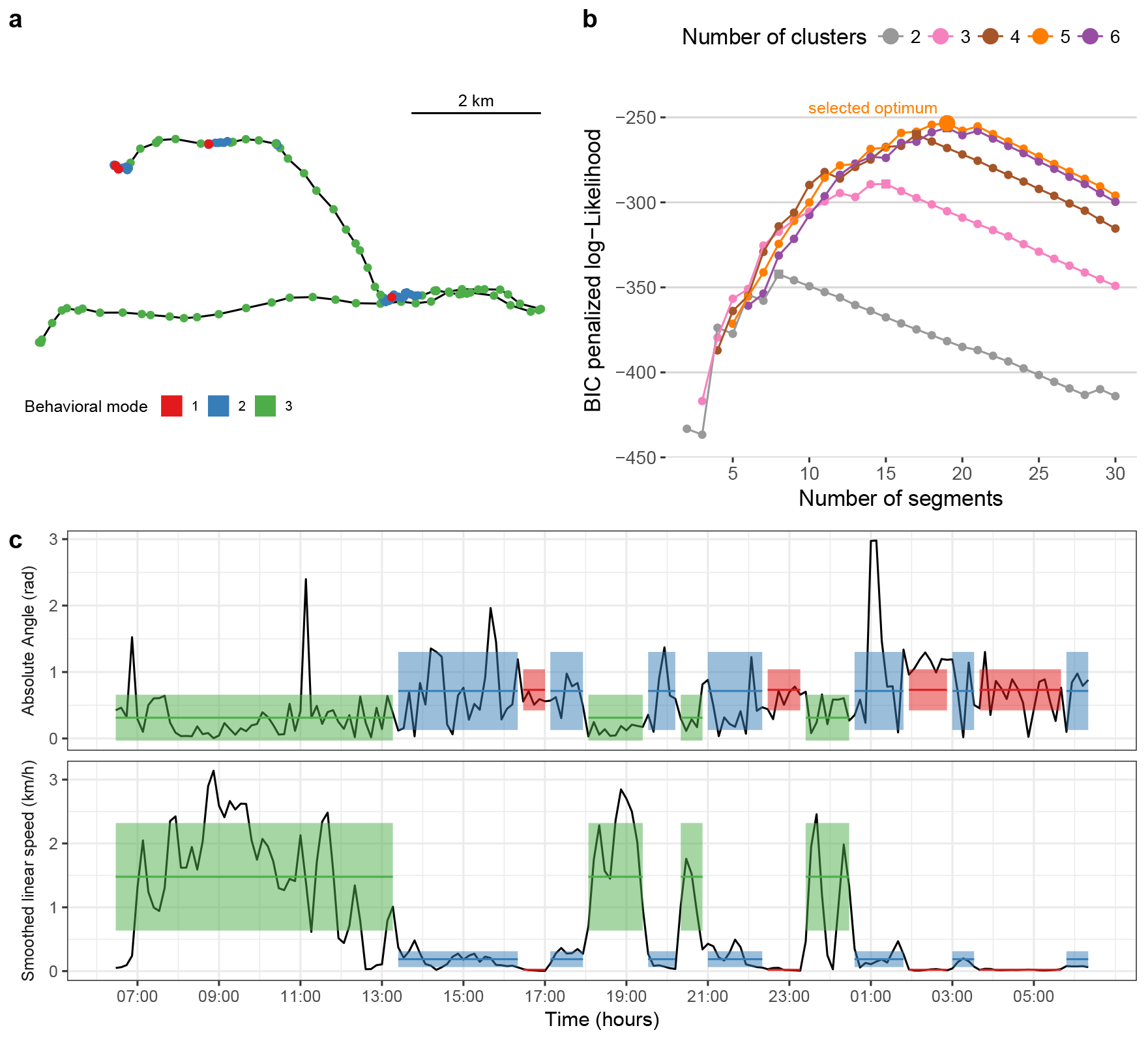
Example of segmentation-clustering of a 24-h zebra’s movement using segclust2d to highlight behavioural changes. (a) Path representation obtained by linking GPS locations recorded every 8 minutes; (b) Determination using BIC-based penalised likelihood of the most likely numbers of states (M = 5) and segments (K = 20) (big orange dot), and most likely numbers of segments for the other number of states considered (large squares at the top of the curves), with Lmin = 5 (i.e. 40 min.). (c) Corresponding time-series in terms of absolute turning angle computed with a constant step length, —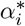—, and smoothed speed, 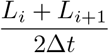, segmented with M = 3 (leading to K = 15); the coloured bands appearing over the two time-series show the estimated occurrences and mean (plain horizontal line running in the middle of the band) +/- standard deviation (band width) for each of the three clusters considered.

#### Illustrative example

We used a 24-h GPS track of a plains zebra (*Equus quagga*) to illustrate the way the segclust2d/segmentation-clustering procedure can identify the occurrences of the various movement modes (Fig. 5). Although that, in this example, the most likely number of modes was estimated to be five, we present the segmentation obtained when setting this number to three, assuming that the biologically relevant modes should be resting (or any other non-moving behaviour such as standing), feeding and transiting (the other two modes detected by our procedure when using five modes were assumed to correspond to mixed behaviours).

**Figure 5:**
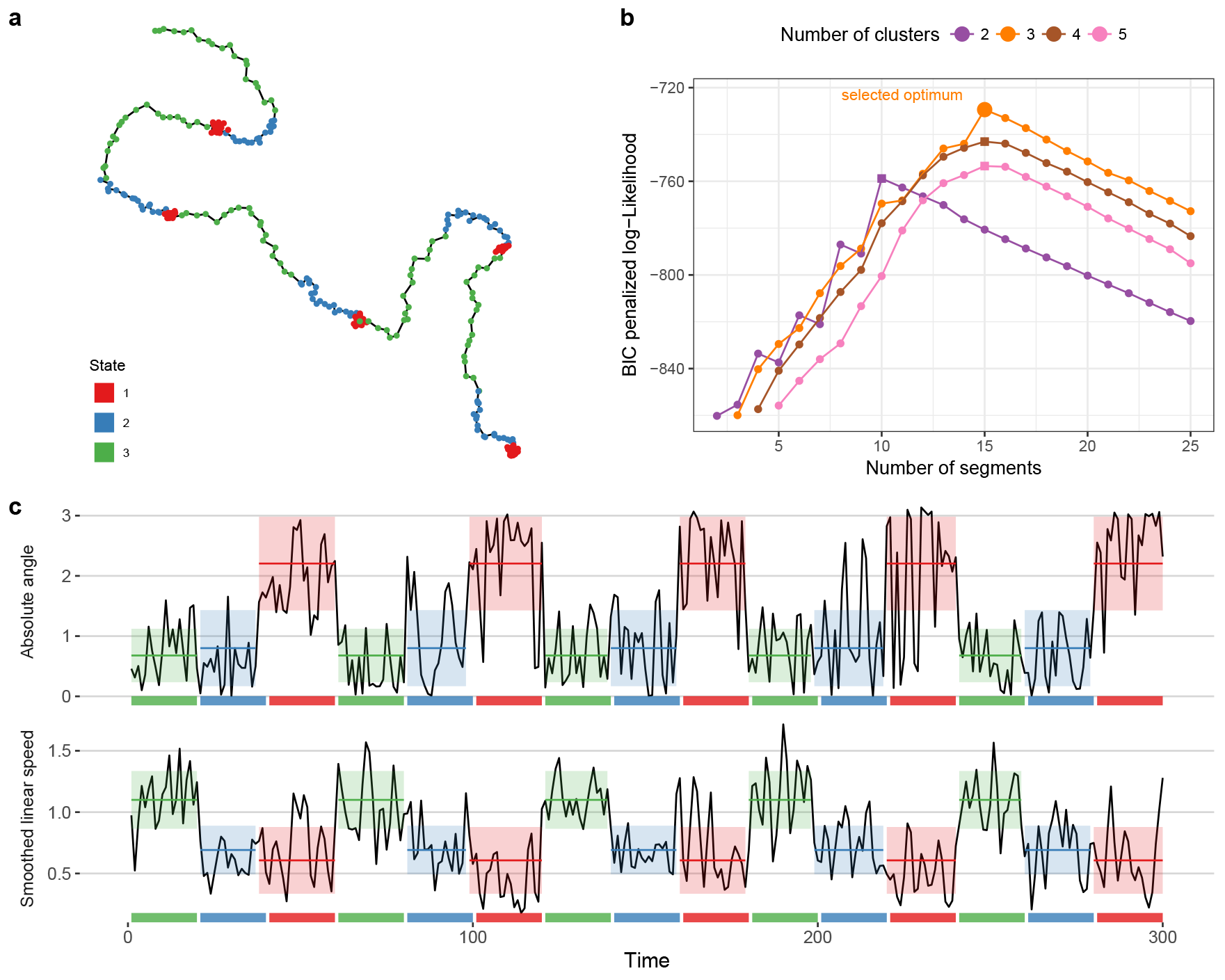
Example of segmentation-clustering of a simulated movement using segclust2d to highlight behavioural changes. (a) Simulated path as a composite correlated random walk, with additional noise *ζ* = 0.3 u; (b) Determination using BIC-based penalised likelihood of the most likely numbers of states (M = 3) and segments (K = 15) (big orange dot), and of the most likely number of segments for the other three numbers of states considered (large squares at the top of the curves). (c) Corresponding time-series in terms of absolute turning angle computed with a constant step length, —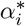—, and smoothed speed, 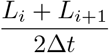, segmented with *L_min_*= 10 and M = 3; the coloured bands appearing over the two time-series show the est +/- standard deviation (band width) for each of the three movement modes whereas the horizontal colour bars running along the time axis show the true occurrences of these modes.

## 4 Discussion

We showed that a generic method, segclust2d,that extends Lavielle (2005)’s and Picard et al. (2007)’s methods to multivariate time-series makes it possible to reliably detect two types of changes that are of key importance when studying free-ranging animal movements: home range shifts, based on bivariate time-series of location coordinates (segmentation-only procedure), and changes in behavioural modes, based on bivariate time-series of turning angles and speed (segmentation-clustering procedure). In any case, this new method is straightforward to parameterize: the user has just to set the minimum segment length (*L_m_in*) to a biologically relevant value so as to prevent the time-series from being over-segmented. Nevertheless, it proved to work at least as well as, and often better than, other recent methods specifically designed to deal with either home range shifts (Gurarie et al., 2017) or changes in behavioural modes (Michelot et al., 2016; McClintock and Michelot, 2018)

Breed et al. (2017) and Gurarie et al. (2017) independently developed an OUM-based method to identify home range shifts in mean location. Using computer simulations, we compared this approach, as implemented in Gurarie et al.’s “marcher” algorithm, with segclutst2d/segmentation-only. Both Gurarie et al. (2017)’s and our method are well able to detect large shifts in mean location. However, our method is also able to detect small shifts in mean location, whereas Gurarie et al. (2017)’s one requires a priori information on the actual mean locations and the shifts dates to correctly detect them, although this is precisely in this case that such information is usually lacking (i.e., they can hardly be guesstimated from visual inspection of the data). Furthermore, contrary to Gurarie et al. (2017)’s method, our method can work with any number of shifts, and is furthermore able to correctly estimate the number of shifts by itself in most cases, and it is also able to reveal changes in home range size. Yet, to be efficient, it does not require any more or less informative initial guess. It simply requires specifying a minimum length (*L_m_in*) for stationary phase to be called a temporary home range, shorter phases being assumed to correspond to transitory exploitations of restricted areas rather than to home ranges. However, contrarily to our method, which considers migrations as simple breakpoints, Gurarie et al. (2017)’s method can estimate the duration of migrations.

The elephant we considered in our illustrative example tended to move back and forth between two main areas. This kind of space use is common in migrating birds that commute between reproductive and wintering home ranges. However, there are numerous studies showing more complex patterns, with an animal setting several distinct temporary home ranges successively (Naidoo et al., 2012; Benhamou, 2014; Cagnacci et al., 2016; Couriot et al., 2018). The segmentation of a long piecewise locational time-series in phases corresponding to temporary home ranges opens promising perspectives to understand how the occurrences and durations of home ranges are related to environmental co-variates, which is a prerequisite to infer long-term consequences for population distribution (Mueller and Fagan, 2008). The elephant illustrative example also shows that, although the model underlying segclust2d looks for stationary phases, there is no guaranty that all segments obtained are really stationary. This occurs because changes between stationary phases are modelled as breakpoints but may in fact correspond to slow progressive changes.

Since the pioneering paper by Morales et al. (2004), HMM-based methods have often been considered the best way to detect changes in behavioural modes of remotely tracked animals. An alternative approach was proposed by Barraquand and Benhamou (2008). It consisted in computing the series of residence time within a virtual circle running along the path and to search for the most likely breakpoints using Lavielle (2005)’s univariate segmentation method. However, although the residence time provides a simple and reliable univariate signal easy to segment and interpret, the values obtained depend not only on the type of behaviour that is performed but also on how long it is performed, preventing the segments corresponding to the same behaviour from being easily clustered. In the present study, we show that the segclust2/segmentation-clustering procedure rivals a HMM-based method when applied to the bivariate signal provided by the two classical metrics that are linear and angular speeds. Importantly, contrarily to what occurs with HMM-based methods, such performance is reached without the need to specify any informative initial state-dependent probability distribution parameters. With both types of method, better results were obtained when using other metrics such as smoothed speed and turning angle measured at constant step length, which were expected to improve the contrast between the intensive and extensive searching modes. Interestingly, using absolute rather than signed values of turning angles measured at constant step length works at best with our method whereas right and left turns were balanced in any mode in our simulated movements. Such metrics should be particularly useful to distinguish between intensive and extensive modes when the former involves turning systematically right or left, i.e. characterized by markedly either negative or positive mean turning angles, whereas the latter involves balanced turning, as occurs in some species (e.g. Smith, 1974). As it results also in a reliable identification when turns are balanced in both intensive and extensive searching, we recommend using it systematically when using our method to distinguish between extensive and intensive searching phases.

In the illustrative example on zebra’s movements, five behavioural modes were detected by the segclust2d/segmentation-clustering procedure in the time-series of smoothed speed and absolute value of the turning angles measured at constant step length. Nevertheless, based on behavioural observations, we chose to segment the time-series with only three modes assumed to correspond to immobility, feeding and transit. Indeed, although our method can estimate the number of states on a statistical basis as the most likely number of clusters, this estimate was not correct for a number of computer simulations whereas behavioural modes were clearly defined. With actual data, there can be some mixing between modes, for instance transit and opportunistic feeding at some times, so that the estimation of the number of relevant modes may become unreliable. Thus, we recommend using the capacity of the segclust2d/ segmentation-clustering procedure to estimate the number of states only when this number cannot be fixed a priori based on biological arguments. A similar conclusion was reached by (Pohle et al., 2017) for HMM-based methods. It is also worth noting that feeding and resting can be distinguished based on movement characteristics only in animals which have to move significantly (with respect to the location recording noise) to feed. For animals which feed mainly without markedly moving, such as some browser herbivores and carcass-eating carnivores, ancillary activity data provided by on-board accelerometers are absolutely required to distinguish these two behavioural modes. As our method can work conjointly on any number of time-series of any nature, future implementation could integrate activity (accelerometer-based) time-series for a better identification of resting vs. active phases.

The segmentation of piecewise stationary time-series, possibly complemented by the clustering of the resulting segments into functional classes, is often key to understanding the dynamics of processes. Based on bivariate time-series of metrics such as location coordinates (northing and easting) or speed and turning angles, segclust2d has the potential to facilitate discovery in the field of movement ecology. If necessary, this approach can apply similarly to three and more dimensions, so as to consider ancillary variables such as activity, as well as other metrics such as distance to a nest, proxies of habitat quality, or any other variable that may be relevant when studying animal movements. Finally, as it can deal with two or more variables of any nature, our approach should be useful not only in movement ecology but also in many other fields.

## Acknowledgement

We thank Clément Calenge for interesting discussions about segmentation issues. The study was partially funded by the grant ANR-16-CE02-0001-01 of the French ‘Agence Nationale de la Recherche’, and the Zone Atelier program of the CNRS. Computer simulations were programed in Pascal and ran thanks to the FreePascal compiler (www.freepascal.org).

## Data accessibility

The code of the method is already publicly available as an R package (https://cran.r-project.org/package=segclust2d). There are only a few data used as examples, and they will be made publicly available as well.

## Supporting Information 1: Complements about segclust2d

### Expectation-Maximization algorithm and smoothing procedure

The Expectation-Maximization (EM) algorithm used to estimate the distribution parameters in the segmentation–clustering model is known to be sensitive to initialisation, so that it may converge to local maxima of the likelihood. This behaviour has some consequences on the parameters estimates but also makes the choice of the number of segments or states complicated. The classical initialisation solution consists in running the algorithm numerous times and just choose the point with the highest value of the log-likelihood, but this strategy is too computationally demanding. To minimize the risk of reaching local maxima within an unacceptable computation time, we propose the following initialisation strategy: (1) perform a pure segmentation of the signal and (2) use a hierarchical cluster algorithm, based on the log-likelihood ratio distance to assign segments to states.

Even with smart initialisation points, however, the EM algorithm may still converge to local maxima. This situation appears when looking, for a given number of states *M*, at the log-likelihood as a function of the number of segments *K*: whereas it is expected to be somewhat regular, this function can be quite noisy. To solve this issue, we propose to ‘smooth’ the function based on the parameter estimates of the distribution (*π_m_*, *μ_m_*, Σ_*m*_) obtained for all the ‘reliable’ solutions, which correspond to the points that lie on the convex hull of log-likelihood curve, as smart initial points for any ‘non-reliable’ solution, i.e. for which an initialisation problem can be suspected. The improvement in terms of regularity of the log-likelihood curve obtained thanks to this procedure is illustrated in figure 6.

**Figure 6:**
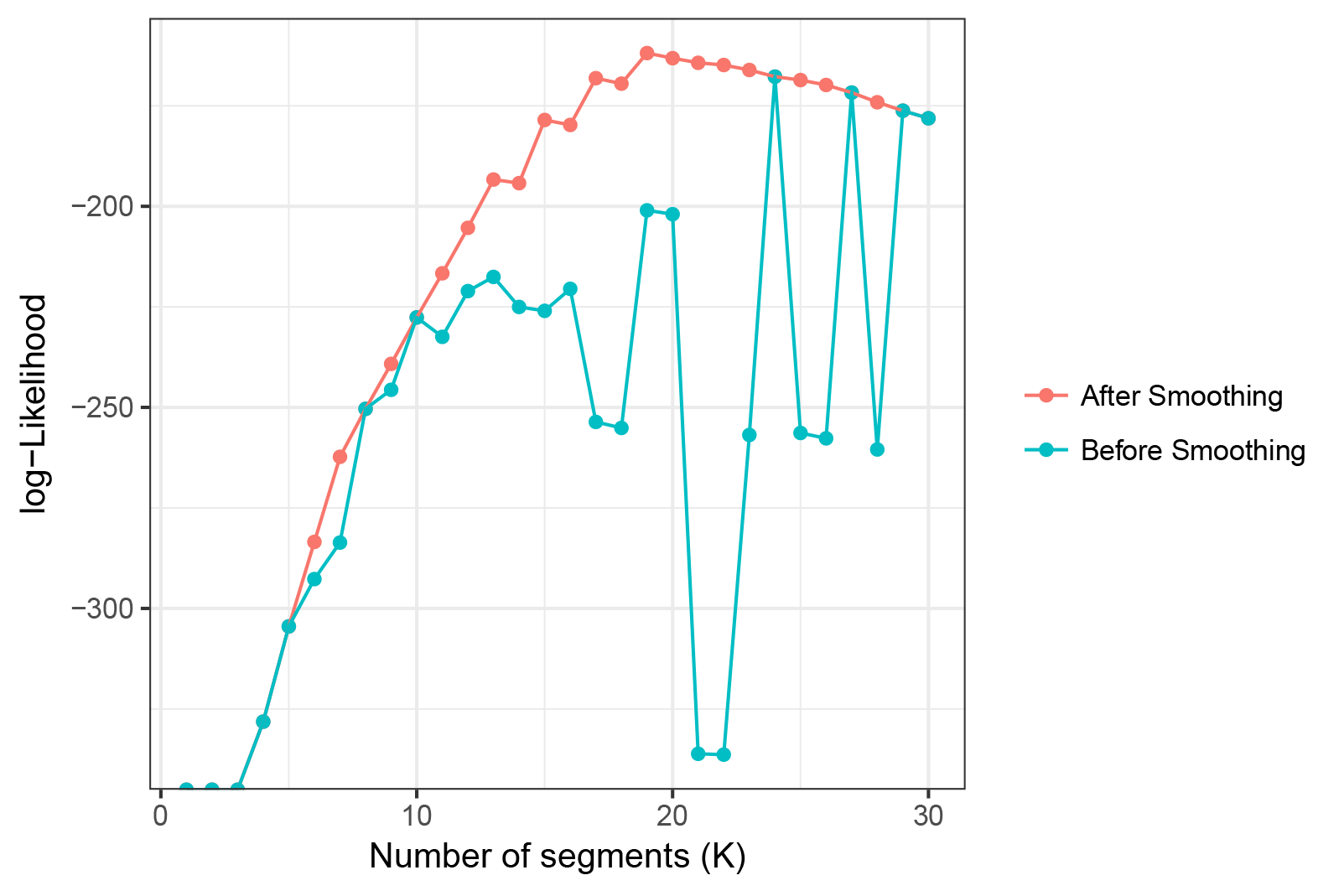
Maximum Likelihood estimates as a function of the number of segments before (in blue) and after (in red) smoothing. For instance the ‘reliable’ solution obtained with *K* = 24 segments was used to provide starting points for the EM algorithm for *K* = 23 and for *K* = 25, and this smoothing procedure gradually spreads over adjacent points.

**Figure 7:**
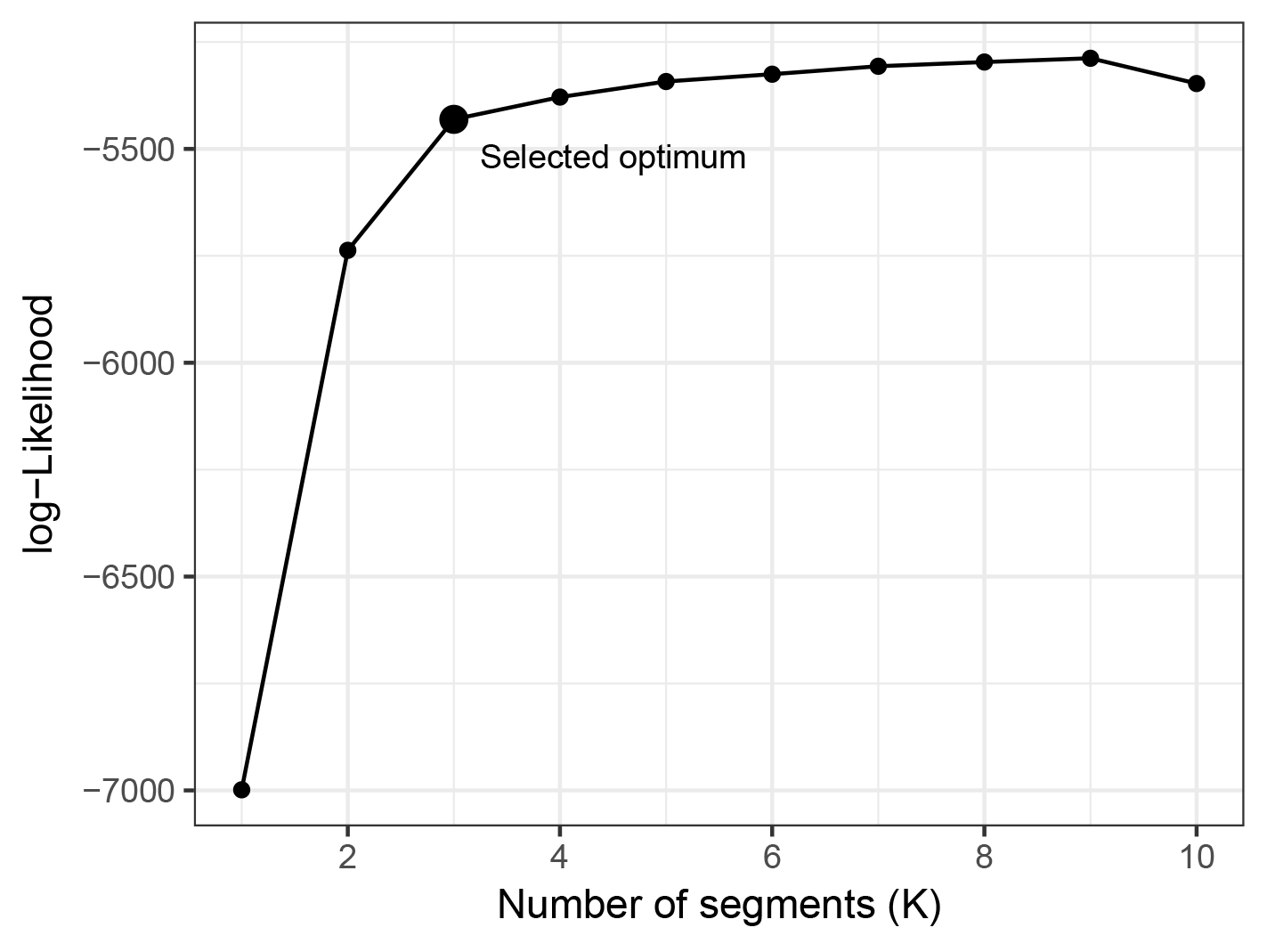
log-likelihood of a segmentation as a function of the number of segment. The optimum selected by the criterion from Lavielle (2005) should be located at a break in the increase of the curve.

### Model selection

#### Choice of the number of segments K in the pure segmentation model

We used the adaptive model selection strategy proposed in Lavielle (2005) consisting in choosing the value of *K* that maximizes the following penalized log-likelihood: ℒ*_K_* − C K where ℒ_*K*_ is the log-likelihood of the optimal segmentation in *K* segments and *C* is a unknown positive constant to be calibrated. The heuristic proposed by Lavielle (2005) consists in detecting the value of *K* for which the log-likelihood ceases to increase significantly. More specifically, consider the normalised log-likelihood defined as

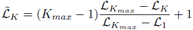

Then, *K* is chosen as the value such that ℒ̃_*K*_ display the largest slope change. Namely, we take:

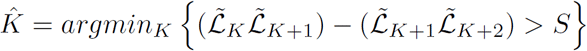

where the value of threshold S is set to a predefined value (we used *S* = 0.7 as proposed in Lavielle, 2005).

As the selection relies on a predefined threshold, it is worth checking where the point corresponding to the selected number of segment lies on a plot of the log-likelihood curve (fig. S3.2). The optimal K value obtained in this way should correspond to a noticeable slope change.

#### Choice of the number of segments *K* and states *M* in the segmentation–clustering model

Selection of the best segmentation-clustering model (i.e. of the best couple of *K* and *M* values) is a hard task as no method has been yet proposed for this purpose. A log-likelihood is expected to increase with the number of parameters. However, as explained in Picard et al. (2007), if the log-likelihood increases with the number of clusters *M*, it does not always increases with the number of segments *K*. Indeed a phenomenon of self-penalization occurs at the ‘true’ number of segments when the detection of breakpoints is easy, stressing to choose *K* simply as the value that maximizes the log-likelihood. However when the detection of breakpoints is more difficult, choosing the maximum value would tend to overestimate *K*. Picard et al. (2007) suggested to add a penalty. A Bayesian Information Criterion (BIC)-based penalty appeared to be sufficient in this case, although it does not work in pure segmentation (Picard et al., 2005). As BIC is the most popular criterion to choose the optimal number of clusters in a mixture model (Frühwirth-Schnatter, 2006), we propose to use the maximum value of the following BIC-based penalised likelihood *B_K,M_* for the selection of both *K* and *M* parameters:

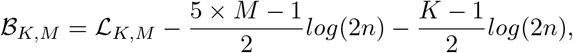

where ℒ_*K,M*_ stands for the log-likelihood for the optimal segmentation-clustering with *K* segments classified into *M* states. The penalization terms in the BIC criterion is half the number of parameters times the logarithm of the size of the dataset. For our model the number of parameters to be estimated is 2*M* means + 2*M* variances + *M* − 1 proportions for the states, and *K* − 1 breakpoints for the segments, and the size of the dataset for *n* bivariate values is 2*n*.

Although this procedure appears to work well for choosing the optimal number of segments, it has been observed to be less reliable for choosing the optimal number of states, which tends to be overestimated. We therefore advise users to set an a priori number of states *M*, based on biological knowledge. We also advise to look at the plot of the BIC-penalized log-likelihood, as in figure 4b of main text, to check that the solution obtained makes sense.

## Supporting Information 2: Interpolating entrance and exit points of a circle

**Figure S2.1:**
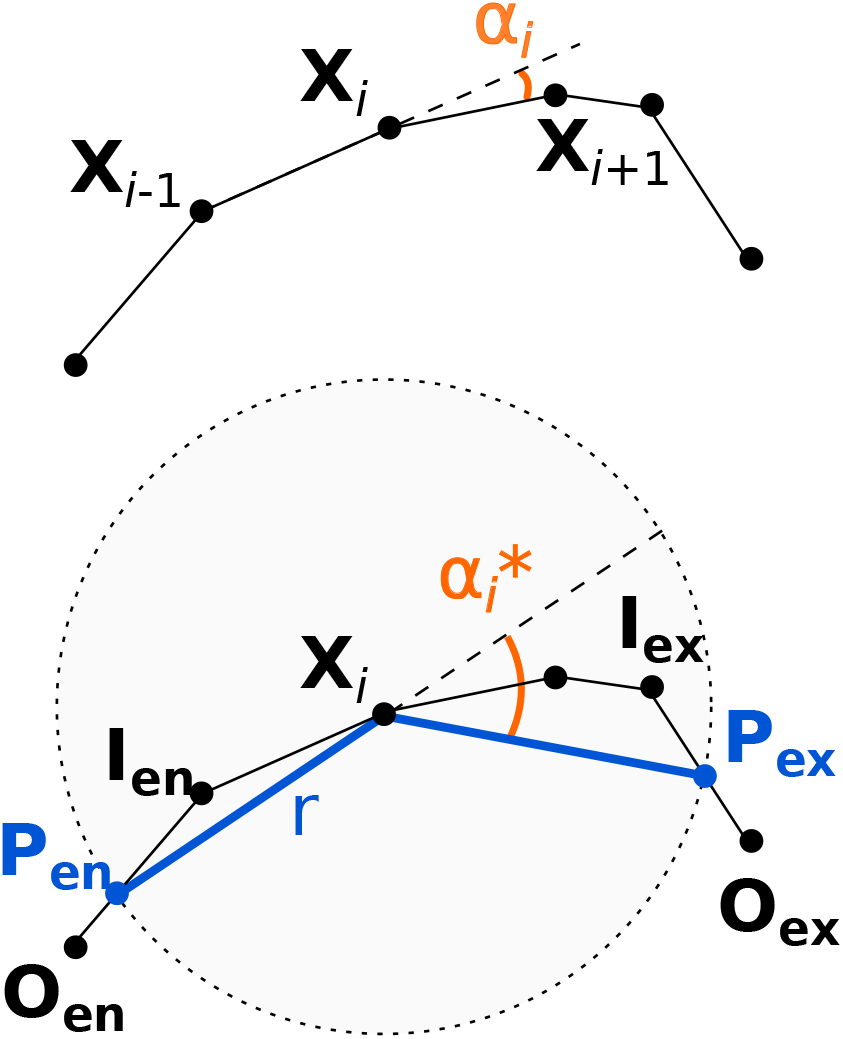
Interpolating entrance and exit points of a circle.

Given a series of locations ***X***_*i*_ = (*x_i_*, *y_i_*) recorded at constant time intervals ∆*t*. Whereas turning angle at constant time intervals *α_i_* (top) corresponds to the change in direction between vectors ***X***_*i−*1_ → ***X***_*i*_ (with length *L_i_*) and ***X***_*i*_ → ***X***_*i*+1_ (with length *L_i_*_+1_) and therefore act as a proxy for angular speeds (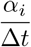), turning angle at constant length interval 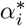 (bottom) corresponds to the change in direction between vectors ***P_en_*** → ***X***_*i*_ and ***X***_*i*_ → ***P_ex_***, where ***P_en_*** and ***P_ex_*** are the last entrance and first exit locations, respectively, of a virtual circle with radius *r* and centred on current location ***X***_*i*_. Let ***I*** = (*x_in_, y_in_*) and ***O*** = (*x_out_, y_out_*) be the last inside and first outside recorded locations, respectively, of the first passage at the circle perimeter, either backwards (***I*** = ***I_en_***, ***O*** = ***O_en_***, ***I_en_*** = ***X***_*i*_ if *L_i_* > *r*) to determine ***P_en_***, or forwards (***I*** = ***I_ex_***, ***O*** = ***O_ex_***, ***I_ex_*** = ***X***_*i*_ if *L_i_*_+1_ > *r*) to determine ***P_ex_***. The location ***P*** (either ***P_en_*** or ***P_ex_***) corresponds to the point where the vector ***I*** → ***O*** intersects the circle perimeter. The length of this vector is 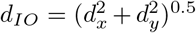, with *d* = *x* − *x*, *d* = *y* − *y*, and its orientation is *θ*, with 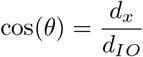 and 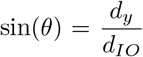. In a new orthonormal frame of reference (*U*, *V*) originating at ***I*** and with *U* axis running through ***O***, the coordinates of current location ***X***_*i*_ become *u_i_* = (*x_i_* − *x_in_*)cos(*θ*) + (*y_i_* − *y_i_n*)sin(*θ*) and *v_i_* == (*y_i_* − *y_in_*)cos(*θ*) + (*x_i_* − *x_i_n*)sin(*θ*). By applying Pythagoras’ theorem, one gets 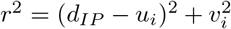, where *d_IP_* corresponds to the distance between ***I*** and ***P***, with *d_IP_* > *u_i_*. Entrance or exit location can therefore be linearly interpolated as 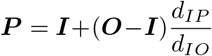 (i.e. *x_P_* = *x_in_* +cos(*θ*)*d_IP_* and *y_P_*= *y_in_*+sin(*θ*)*d_IP_*), with 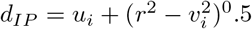.

## Supporting Information 3: Efficiency of the segmentation-clustering method

### 1 Comparison with HMM for low (*ζ* = 0.2) and high (*ζ* = 0.4) noise levels

**Figure S3.1:**
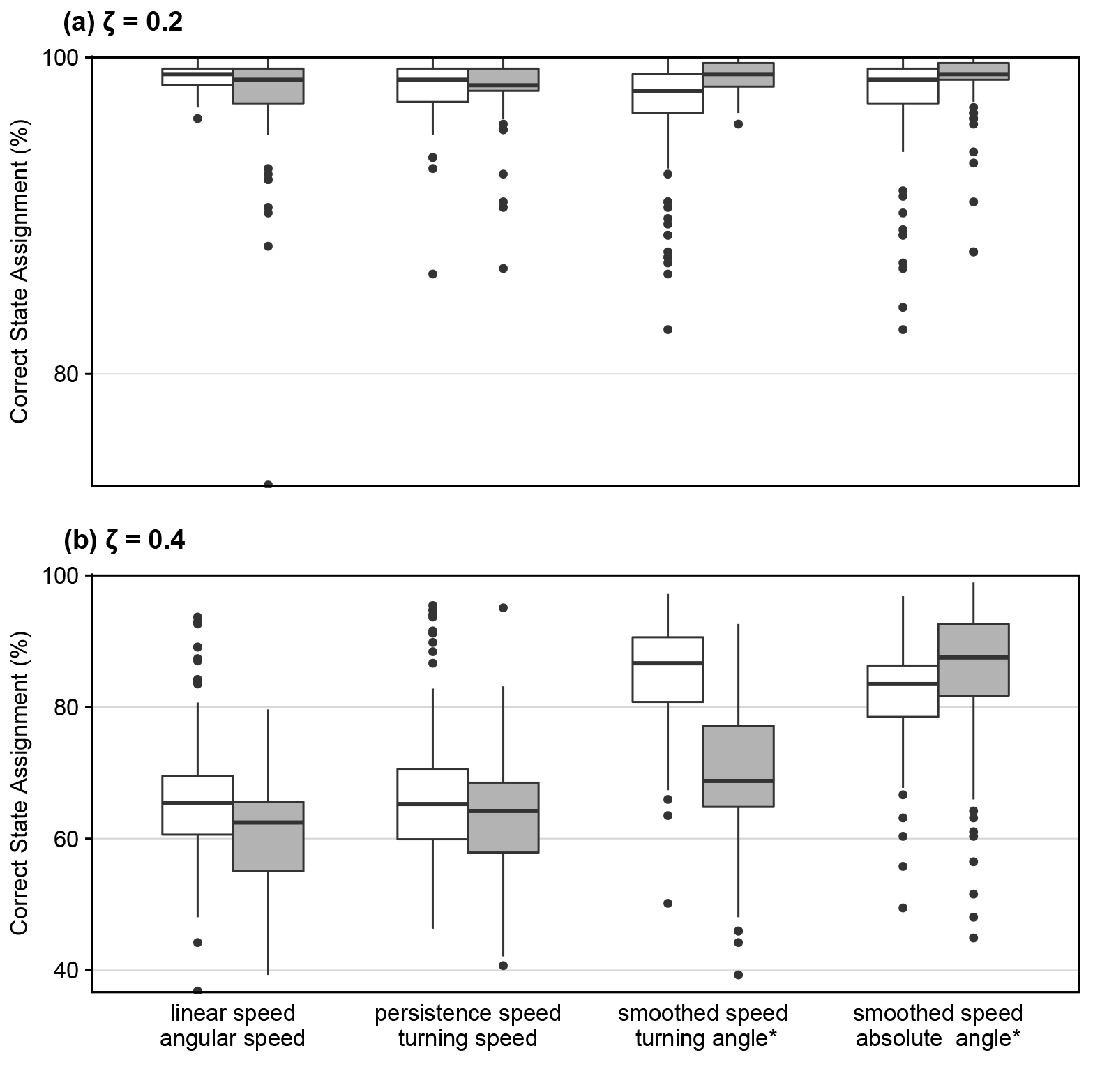
The boxplots show the proportion of correct state assignments, obtained for various bivariate signals when the true number of states is known (M = 3) with noise level *ζ* = 0.2*u* (a) or *ζ* = 0.4*u* (b), as estimated from 100 replicates. The star (*) indicates turning angles computed with a constant step length, in terms of arithmetic (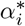) or absolute (—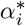—) values. The white boxplots show the results obtained with HMM-based R package momentuHMM (McClintock and Michelot, 2018), with informative initial state-dependent probability distribution parameters set to the true values of the various metrics in the different states (using the following distributions: Gaussian for persistence and turning speeds, wrapped Cauchy for angular speed and turning angle 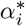, Weibull for linear speed, smooth speed and —turning angle*—). The grey boxplots shows the results obtained using the segclust2d/segmentation-clustering procedure with *L_min_* = 10.

### 2 Estimation of the number of states

**Figure S3.2:**
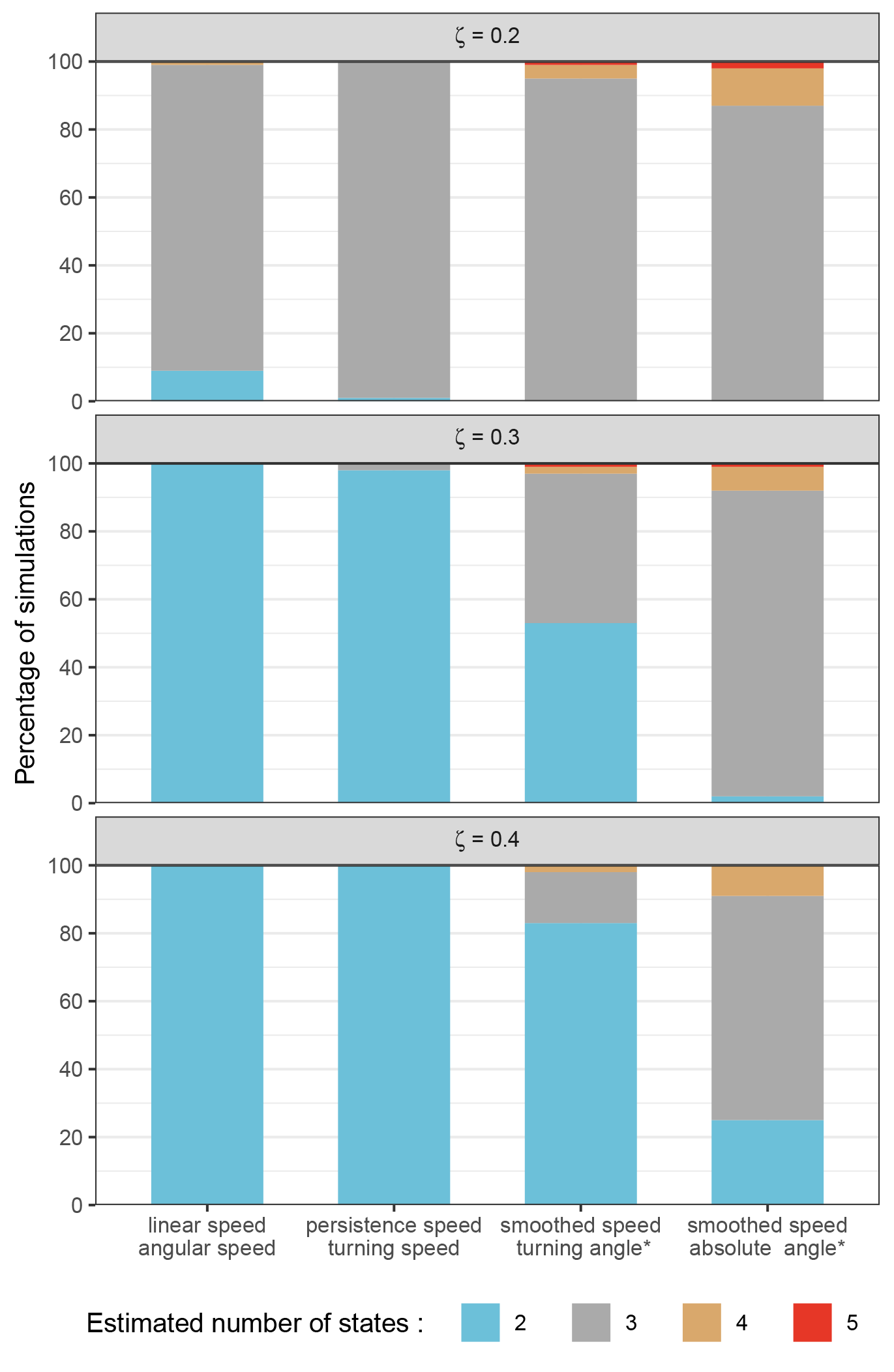
The various bars show the proportions of simulations resulting in a predicted number of states (i.e. behavioural modes) equal to 2, 3, 4, or 5, for the three noise levels considered (*ζ* = 0.2*u*, *ζ* = 0.3*u* and *ζ* = 0.4*u*) and the four types of couples of metrics considered. The true number of states is 3. The star (*) indicates turning angles computed with a constant step length, in terms of arithmetic (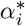) or absolute (—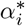—) values. The couple of metrics leading to best segmentation when the true number of states is known – absolute turning angle computed with a constant step length and smoothed speed – also leads to the best estimation of the number of states, but this latter estimation is not fully satisfactory, and should be worse with actual data because of possible mixing of movement behaviours.

**Authors’ contributions.** RP analysed the data and contributed to the coding of the statistical model, which was developed by MPE and EL. SC provided the tracking data. SB led the project, performed computer simulations, and wrote the first draft of the manuscript, except the part describing the model which was first contributed to by MPE, EL and RP. All authors contributed significantly to the final manuscript.

